# Interleaved single and bursting spiking resonance in neurons

**DOI:** 10.1101/2024.06.24.600479

**Authors:** Cesar C. Ceballos, Nourdin Chadly, Eric Lowet, Rodrigo F. O. Pena

## Abstract

Under *in vivo* conditions, CA1 pyramidal cells from the hippocampus display transitions from single spikes to bursts. It is believed that subthreshold hyperpolarization and depolarization, also known as down and up-states, play a pivotal role in these transitions. Nevertheless, a central impediment to correlating suprathreshold (spiking) and subthreshold activity has been the technical difficulties of this type of recordings, even with widely used calcium imaging or multielectrode recordings. Recent work using voltage imaging with genetically encoded voltage indicators has been able to correlate spiking patterns with subthreshold activity in a variety of CA1 neurons, and recent computational models have been able to capture these transitions. In this work, we used a computational model of a CA1 pyramidal cell to investigate the role of intrinsic conductances and oscillatory patterns in generating down and up-states and their modulation in the transition from single spiking to bursting. Specifically, the emergence of distinct spiking resonances between these two spiking modes that share the same voltage traces in the presence of theta or gamma oscillatory inputs, a phenomenon we call interleaved single and bursting spiking resonance. We noticed that these resonances do not necessarily overlap in frequency or amplitude, underscoring their relevance for providing flexibility to neural processing. We studied the conductance values of three current types that are thought to be critical for the bursting behavior: persistent sodium current (*I*_NaP_) and its conductance *G*_NaP_, delayed rectifier potassium (*I*_KDR_) and its conductance *G*_KDR_, and hyperpolarization-activated current (*I*_h_) and its conductance *G*_h_. We conclude that the intricate interplay of ionic currents significantly influences the neuronal firing patterns, transitioning from single to burst firing during sustained depolarization. Specifically, the intermediate levels of *G*_NaP_ and *G*_KDR_ facilitate spiking resonance at gamma frequency inputs. The resonance characteristics vary between single and burst firing modes, each displaying distinct amplitudes and resonant frequencies. Furthermore, low *G*_NaP_ and high *G*_KDR_ values lock bursting to theta frequencies, while high *G*_NaP_ and low *G*_KDR_ values lock single spiking to gamma frequencies. Lastly, the duration of quiet intervals plays a crucial role in determining the likelihood of transitioning to either bursting or single spiking modes. We confirmed that the same features were present in previously recorded in vivo voltage-imaging data. Understanding these dynamics provides valuable insights into the fundamental mechanisms underlying neuronal excitability under *in vivo* conditions.

**Author summary:** Since discovering that neurons in the hippocampus can encode spatial position through phase precession, many experiments have explored how specific theta and gamma oscillations influence location specificity in the brain. However, the individual neuronal properties and dynamics behind these behaviors are still being uncovered. Previously, we found that stereotypical bursting and single-spike firing in pyramidal neurons are linked to these oscillations and further associated with an animal entering or leaving a place field. Advances in voltage-imaging techniques have enabled us to assess these properties more precisely. Our study shows that different frequencies can independently trigger these stereotypical spikes, demonstrating a complex pattern where the same cell can be double-coded: a phenomenon we called interleaved resonance. Additionally, we found that this coding can be modulated by persistent sodium and delayed-rectifier potassium currents. Moreover, these neurons are more likely to burst following long periods of silence. These findings provide new insights into the mechanisms underlying neural coding in the hippocampus and how it relates to behavior.

## Introduction

Neurons from the CA1 hippocampus display a diversity of spiking dynamics. In the hippocampus, from the point of view of individual cells, *in vivo* voltage imaging [1, 2, 3], patch clamp recordings [4, 5, 6, 7], and multielectrode recordings [8, 9] reveals that CA1 pyramidal cells exhibit stereotypical single and burst spiking. While investigating their emergence and transitions it was noted that these spike modes appear with different probabilities of occurrence where bursts are more likely to occur after long periods of activity silence [9]. From the point of view of populations, gamma (30-100 Hz) and theta (3-12 Hz) oscillations coexist and have been well characterized in CA1 region of rodent hippocampus [10, 11]. Gamma is modulated by theta during working memory retention, and it is important to notice the cross-frequency coupling of these oscillations for spatial navigation and memory performance [12, 13]. Recent studies have tried to connect those transitions using *in vivo* data, as shown by shifts from single to bursting spikes in the same neurons that are preferentially observed during theta and gamma oscillations, respectively [1]. Here we use the term burst to define cluster of spikes with intervals lower than 14 ms, according to studies where this variation is found ∼6-14 ms [9, 4, 14]. Behaviorally, these spiking modes have been proposed to encode an animal’s position, where burst signals when animals enter a place. In contrast, single spikes signal when the animal leaves a place [1].

Admittedly, understanding spiking patterns transitions is synonymous to understanding ionic current’s dynamics. For fixed synaptic input, the composition of ionic currents can produce distinguishable spiking outputs that vary from single to burst of spikes [15, 16, 17]. Combinations of ionic currents can determine these spike modes as in the case where expression of calcium is linked to emergence of burst [18, 19, 20], or in combination with the spatial location on the neuron as in the case of persistent sodium current (*I*_NaP_) located at or near the soma [15, 21], or only the geometry [22], or combinations of both [23]. Moreover, potassium currents such as *I*_h_ have been investigated in terms of their effects on subthreshold oscillations and resonance [24, 25, 26, 27], although their influence on bursting has still not been clarified (but see [28] for modeling efforts on its influence on this direction).

Oscillations can cause transitions from low to high firing rate on a neuron. Cells may have amplified response for certain oscillatory frequencies compared to other frequencies. This is known as suprathreshold (spiking) resonance, and it has been observed widely across the brain [29, 30, 31, 32, 33], including CA1 pyramidal cells [34] and the mechanisms underlying this resonance have been intensively investigated using computational models [35, 36]. Similarly, subthreshold resonance can be seen as a transition voltage change of a neuron upon a particular (non-zero) input frequency [37, 38]. Several ionic currents contribute to subthreshold resonance and can be discussed in terms of their chord and derivative conductances [39]. For example, persistent sodium current (*I*_NaP_) is an amplifying ionic current associated to bursting in CA1 [40] and delayed rectifier potassium (*I*_KDR_) is a resonant current [37, 41, 42]. Less is known about the role of these ionic currents on suprathreshold (spiking) resonance.

It is only natural to assume that CA1 pyramidal neurons embedded in a network can take advantage of the spiking resonance to preferentially transmit information encoded in the spiking mode (single or burst spiking) when a particular oscillatory synaptic input (e.g. theta or gamma oscillations) is received [34, 29, 43, 44]. In addition to that, it is important to understand how transitions at these different scales connect and modulate them. In this paper, we investigate suprathreshold resonance of single and burst spiking in a CA1 pyramidal neuron model. We found that spiking resonance can easily emerge when neurons are stimulated with theta and gamma synaptic inputs and ionic currents are tuned. Variability of sodium and potassium currents differentially promoted the presence of resonance according to the firing pattern (single vs. burst), a phenomenon we call interleaved single and bursting spiking resonance. Our results provide insights that might help to understand the mechanisms underlying single and burst spiking computation and its role in connection to *in vivo* processing that is relevant for spike coding in CA1 pyramidal cells for spatial navigation and memory.

Our paper is organized as follows: in the following section, we present studies of how ionic currents promote transitions from single to burst firing under constant sustained input and later under oscillatory inputs. For the oscillatory inputs, we show how resonance emerges and depends on ionic currents. Then we move to a more systematic study where we characterize interleaved single and bursting resonance in the presence of dendritic *I*_h_ current. For the same model, we characterize phase-locking to gamma and theta and how these main ionic currents shape this relationship. We conclude by systematically characterizing the probability of bursting spikes in relation to the periods of silence in the voltage traces. Methods are included in the last section of the paper.

## Results

### Ionic currents promote a transition from single to burst firing during sustained depolarization

CA1 pyramidal cells are known to exhibit transitions from single (SS) to bursting spikes (BS) *in vivo* [1, 2, 3, 45]. This transition occurs in the presence of subthreshold oscillations and is modulated by oscillatory inputs, mostly theta and gamma oscillations. Subthreshold activity is characterized by transitions from hyperpolarized states to depolarized states (also known as down and up-states, respectively). Firing mostly occurs on top of the depolarization phases. Input patterns play a relevant role in the switching, but it is unclear how they can be promoted by the neuron’s intrinsic properties, e.g. ionic currents. We first decided to investigate the firing patterns occurring during depolarization states and how SS and BS emerge under different ionic conductances.

We first investigated the role of conductances that are mostly expressed in the soma, such as *I*_NaP_ and *I*_KDR_. In this section, we used a single compartment computational model of CA1 pyramidal cell (see Methods) and we varied *G*_NaP_ (persistent sodium conductance) and *G*_KDR_ (delayed rectifier potassium conductance) for the model receiving a step function of current that maintains the voltage in a constant depolarized state (Fig 1A). We observed that increasing *G*_NaP_ makes the firing transits from SS to BS and finally to plateau potential, also known as depolarization block [46] (see voltage traces Fig 1B1). The phase plots (dV/dt) in the same figure reveals how the voltage series transits between the different modes (Fig 1B2). In contrast, the effect of increasing *G*_KDR_ is a transition from BS to SS (Fig 1C1) and this can be seen in the phase plots (Fig 1C2). Single spiking can be explained as a result of having long pauses between spikes due to long afterhyperpolarization, which is facilitated by strong action potential repolarization due to large *G*_KDR_ or by preventing fast depolarization due to small *G*_NaP_.

**Fig 1.**
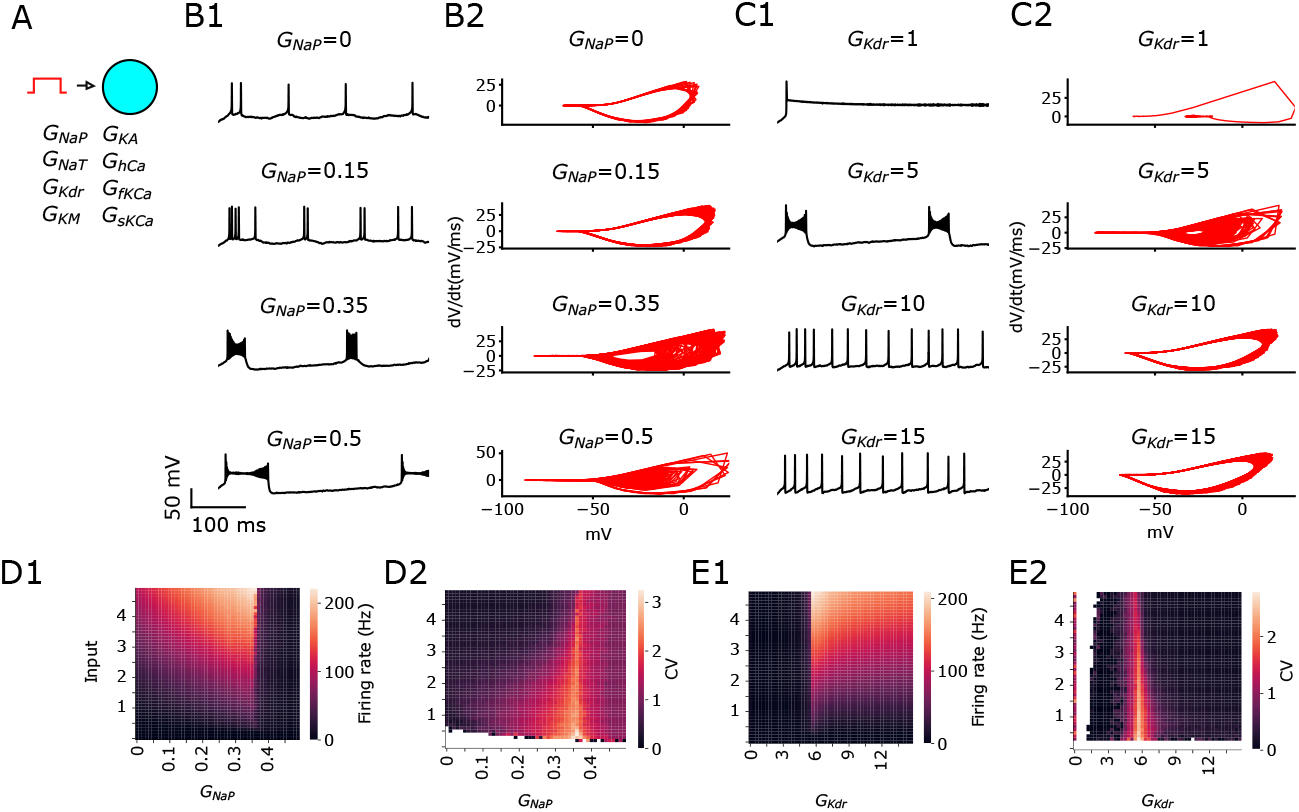
Ionic currents promote transition from SS to BS during sustained depolarization. Default values for fixed conductance while varying the other are *G*_NaP_ = 0.3 and *G*_KDR_ = 6. **A** Sketch of CA1 model receiving a step function of current. **B1** Example voltage traces and **B2** phase plots (dV/dt) when varying *G*_NaP_. **C1** Example voltage traces and **C2** phase plots (dV/dt) when varying *G*_KDR_. **D1** Firing rate and **D2** coefficient of variation (CV), a measure the activity of the neuron and the presence of bursts, displayed as heatmaps spanned over *G*_NaP_. **E1** Firing rate and **E2** coefficient of variation (CV) displayed as heatmaps spanned over *G*_KDR_ vs. Input current. For phase plots we remove the first 100 ms, but for all other panels we show the traces from the beginning with stimulation application starting at 0 ms.

The firing rate increases with the increase of stimulation amplitude, increase of *G*_NaP_ and decrease of *G*_KDR_ (Fig 1D1 and 1E1, respectively). As a measure of the burstiness we adopted the coefficient of variation (CV), as described in the methods. The CV above 2 (reddish to yellowish) determines the area in which the combination of conductance promotes BS behavior (Fig 1D2 and 1E2). As it is shown in the heatmaps, BS is independent of the input amplitude and mostly occurs in areas around *G*_NaP_ = 0.35 or *G*_KDR_ = 6 (Fig 1D2 and 1E2, respectively). These values correspond to the edges where the transition from spiking to depolarization block occurs (see dark regions in heatmaps of firing rate). Interestingly, BS occurs in a very narrow range of conductances, meanwhile single spiking is likely to occur in a wide range of values. This set of observations shows the opposite effects of the *G*_NaP_ and *G*_KDR_ ionic currents in modulating a switch between SS to BS. We have also investigated the effect of the additional ionic currents *I*KA the A-type K^+^, *I*KM the muscarinic-sensitive potassium, *I*sKCa the slow Ca^2+^-activated potassium current, and *I*fKCa the fast Ca^2+^ -activated potassium current (Fig. S1). Only *G*_KM_ caused a significant effect on the modulation of bursting behavior.

### Intermediate *G*_NaP_ and *G*_KDR_ generate spiking resonance preferentially at gamma frequency input

We showed previously that *G*_NaP_ and *G*_KDR_ have opposite effects on the generated spiking patterns during sustained depolarization. Nevertheless, CA1 pyramidal neurons receive oscillatory inputs in the range from 3 to 100 Hz, where oscillatory gamma is between 30-100 Hz and theta 3-12 Hz frequencies. This induces transitions from hyperpolarized to depolarized states where shorter intervals occur at high input frequencies such as gamma frequencies and longer intervals at theta frequencies. Under this condition, firing during depolarization is not sustained, but rather is alternated by periods of silence. In our paper, we investigate the role of the silent states generated by the interplay between input patterns and ionic currents to create a suprathreshold resonance and its relation to SS and BS.

For stimulation of the cell in this section, we choose varying oscillatory stimulation frequencies covering the range from theta to gamma frequencies (Fig 2A). In Figs 2B1, B2, D1, and D2 we present voltage-traces for variations of *G*_NaP_ and *G*_KDR_ under theta or gamma (see captions). We investigate the firing rate responses and their preference for a certain input frequency (i.e. suprathreshold resonance). We observe low-pass filtering of the firing rate for low *G*_NaP_ or high *G*_KDR_, suprathreshold resonance for intermediate *G*_NaP_ or *G*_KDR_, and a nearly absent frequency preference for high values of *G*_NaP_ or low values of *G*_KDR_ (Fig 2C, E). Interestingly, regardless of the opposite effects of *G*_NaP_ and *G*_KDR_, we observed that theta input has a stronger effect on the firing rate response than gamma when the neuron acts as a low-pass filter. In contrast, gamma is stronger than theta for resonance (Fig 2C, E). These results unveiled an interaction between the input frequency and the filtering properties that determine the firing output.

**Fig 2.**
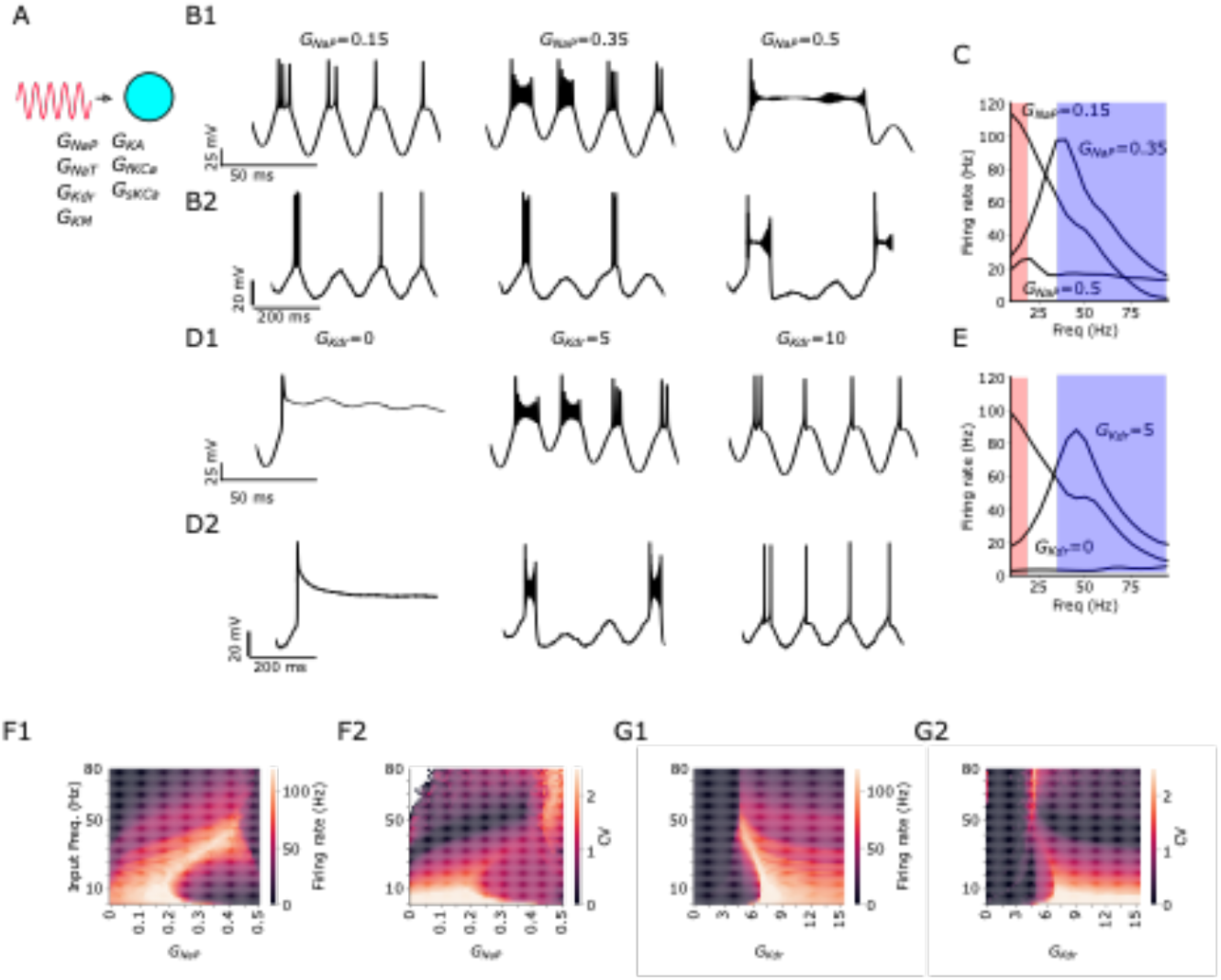
Intermediate *G*_NaP_ and *G*_KDR_ generates firing rate resonance. Default values for fixed conductance while varying the other are *G*_NaP_ = 0.3 and *G*_KDR_ = 6. **A** Sketch of simplified CA1 model receiving oscillatory current stimulation. **B1** Examples of voltage traces for varying values of *G*_NaP_ under gamma stimulation. **B2** Examples of voltage traces for varying values of *G*_NaP_ under theta stimulation. **C** Firing rate while varying stimulation frequency of an oscillatory input. Shaded red area: theta frequencies. Shaded blue area: gamma frequencies. **D1** Examples of voltage traces for varying values of *G*_KDR_ under gamma stimulation. **D2** Examples of voltage traces for varying values of *G*_KDR_ under theta stimulation. **E** Firing rate while varying stimulation frequency of an oscillatory input. **F1** Firing rate and **F2** CV heatmap spanned over *G*_NaP_ and oscillatory input frequency. **G1** Firing rate and **G2** CV heatmap spanned over *G*_KDR_ and oscillatory input frequency. In all panels, stimulation starts at 0 ms. To allow slow but non-zero frequencies to be studied we cut-off the simulations at 2 Hz.

The CVs above 2 (reddish to yellowish) obtained while changing the oscillatory input frequency and the conductance indicate the area where BS occurs (Fig 2F2 and 2G2). The heatmaps show similar patterns for firing frequency and CV for both *G*_NaP_ and *G*_KDR_, i.e. we observe high firing rates simultaneously with high values of CV (Fig 2F1, 2F2, 2G1 and 2G2). The high firing rates obtained can be justified by the BS and low firing rates are by the SS (Fig 2F1, 2G1). Note that for these panels we used a cut-off of 2 Hz to allow slow frequencies to be tested but different from zero. Also note that certain conductance value choices may lead the neuron into non-physiological states, which are most likely to occur at low *G*_KDR_ values and high *G*_NaP_ values which is acceptable for exploratory purposes.

We observed BS at gamma frequencies when *G*_NaP_ levels were intermediate or high (see the yellowish diagonal in Fig. 2F2) and when *G*_KDR_ was at intermediate levels (Fig. 2G2). Additionally, BS was noted at theta frequencies with low *G*_NaP_ or high *G*_KDR_. Similarly, we observed SS at low *G*_KDR_ and at gamma frequencies with low or intermediate *G*_NaP_ or high *G*_KDR_. Interestingly, BS in gamma frequencies occurs simultaneously with resonance at intermediate values of *G*_NaP_ or *G*_KDR_. These results in Figs 2F1 and 2F2 indicate that when suprathreshold resonance is present, a linear increase in input frequency and *G*_NaP_ shifts the BS presence from theta to gamma frequencies. Similarly, a decrease in *G*_KDR_, along with increased input frequencies, also shifts the BS presence from theta to gamma frequencies (as shown by the yellowish regions in the heatmaps, Fig 2G1 and 2G2). A similar pattern is observed for SS, with the dark regions in the *G*_NaP_ heatmap and the dark middle-right region in the *G*_KDR_ heatmap illustrating this shift (Fig 2F2 and 2G2, respectively). We have also investigated the effect of the additional ionic currents *I*KA the A-type K^+^, *I*KM the muscarinic-sensitive potassium, *I*sKCa the slow Ca^2+^-activated potassium current, and *I*fKCa the fast Ca^2+^ -activated potassium current (Fig. S2). Only *G*_KM_ caused a significant effect on the modulation of bursting behavior. However, the effect of *G*_KM_ was restricted to gamma band, meanwhile for *G*_NaP_ and *G*_KDR_ we observed effects in both gamma and theta. Combined with the results in Fig. S1, we therefore decided to focus on the effect of *G*_NaP_ and *G*_KDR_ for the remainder of the paper.

In summary, for low *G*_NaP_ and high *G*_KDR,_ we found high firing rate response to low input frequencies (i.e. theta) due to weak filtering, and low firing rate response to high input frequencies (i.e. gamma) due to strong filtering. This suggests that the tuning of ionic currents in interaction with input frequencies are critical for determining the occurrence of resonance.

### Single and burst firing display suprathreshold resonance with different amplitudes and different resonant frequencies

It is well known that CA1 pyramidal cells have a high expression of the hyperpolarization activated *I*_h_ current [47]. This current is mostly expressed in the dendrites and plays a relevant role in modulating synaptic inputs [47, 48, 49, 50] and resonance [39]. It prevents excessive hyperpolarization and causes rebound spiking after hyperpolarizations [51, 52]. Thus, in this section, we incorporated a dendritic compartment containing *I*_h_ current (Fig 3A). Following previous work [1], *I*_h_ is also necessary for the transition single-spike/bursting behavior and therefore it is added in the next sections.

**Figure 3.**
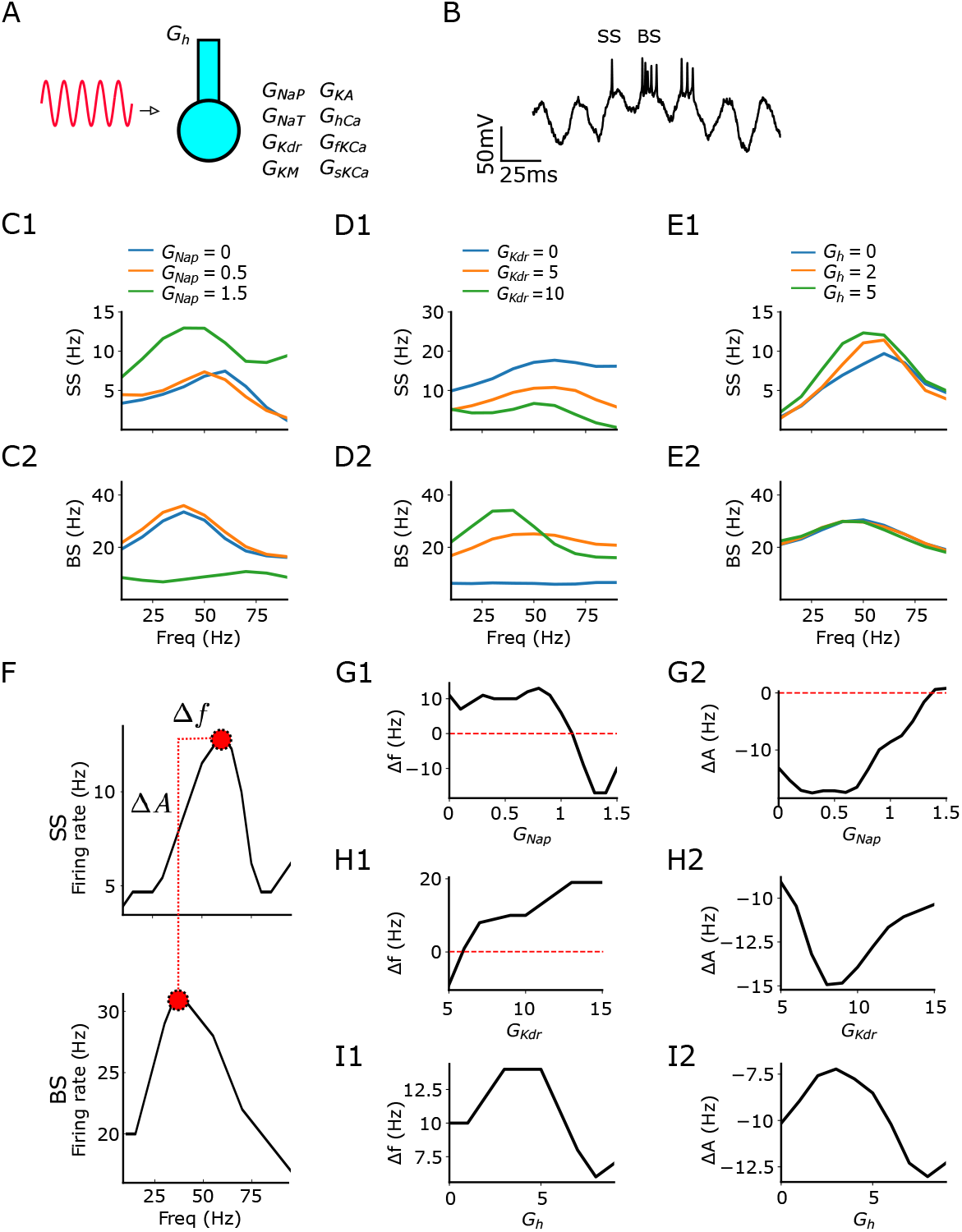
Single and burst firing display suprathreshold resonance with different amplitudes and resonant frequencies. Default values for fixed conductance while varying the other are *G*_naP_ = 0.93, *G*_KDR_ = 5.86, and *G*_h_ = 8.46. **A** Sketch of CA1 model with additional dendritic compartment receiving oscillatory current stimulation. **B** Exemplary voltage trace of the simulation with SS and BS indicated. **C1-2** Firing rate vs oscillatory input frequency for SS **C1** and BS **C2** when *G*_NaP_ values are 0, 0.5 and 1.5. **D1-2** Firing rate vs oscillatory input frequency for SS **D1** and BS **D2** when G_KDR_ values are 0, 5 and 10. **E1-2** Firing rate vs oscillatory input frequency for SS **E1** and BS **E2** when *G*_h_ values are 0, 2 and 5. **F** Representative sketch demonstrating how Δ*f* and Δ*A* are calculated for the interleaved SS and BS resonances. **G1-2** Δ*f* and Δ*A* for varying values of *G*_NaP_. **H1-2** Δ*f* and Δ*A* for varying values of *G*_KDR._ **I1-2** Δ*f* and Δ*A* for varying values of *G*_h_. Colors mark the magnitude of ionic currents (blue = low, orange = medium, green = high).

We stimulated the cell model with varying oscillatory stimulation frequencies covering the range from theta to gamma frequencies (Fig 3A). Under this condition, the neuron is capable of switching its firing from SS to BS (Fig 3B). We separated our quantitative measures into SS and BS (see Methods). Overall, we observed resonance for all the explored conductances and for both spiking types (SS and BS), except for BS when *G*_NaP_ is large or when *G*_KDR_ is low (Fig 3 C2, D2). Finally, SS and BS have resonance for all *G*_h_ values (Fig 3 E1, E2).

We further compared the resonance frequencies and their respective firing rate amplitudes between SS and BS (Fig 3F). For low *G*_NaP_, SS resonance frequency is larger than BS resonance frequency, but it is the opposite when *G*_NaP_ is large (Fig 3G1). For low *G*_KDR_, SS resonance frequency is smaller than BS resonance frequency, but it is the opposite when *G*_KDR_ is large (Fig 3H1). Moreover, SS resonance frequency is larger than BS resonance frequency for all *G*_h_ values (Fig 3I1). Finally, as expected, firing frequency at the resonant peak was always higher for BS rather than SS (Fig 3G2, H2, I2).

The set of results in this section demonstrates an interesting coding behind SS and BS in our CA1 model. First, it shows that neither the resonance frequencies nor resonance amplitudes between SS and BS necessarily overlap. More interestingly, this phenomenon happens at the same voltage traces (interleaved) and can be modulated by ionic currents.

To gain more insight about the mechanism underlying the generation of interleaved single spikes and burst, in particular for currents that were not studied in depth in our analysis, we compared conditions where the *I*_KM_ is present and when it is zeroed, the latter a simulation of the experimental effect of channel blocking. Voltage traces show that decreasing *I*_KM_ affect mostly single spiking, specifically converting them into bursts (Fig S2E; notice first, third, and seventh cycle where single spike become burst). Furthermore, we characterized the time evolution of activation variables of *I*_KDR_, *I*_KM_ and *I*_NaP_ (Fig S3). We observed different time scales, with *I*_NaP_ being the fastest current, *I*_KDR_ being slower and *I*_KM_ the slowest current. This was expected from the way their kinetics was set up (see Methods).

We also investigated the effect of the background noise on the interleaved resonance from single spiking to burst. We found the interleaved resonance is sensitive to the noise level (Fig S4). Altogether, these results suggest that the interleaving from single spiking to burst is a consequence of the interplay of three ionic currents (*I*_KDR_, *I*_KM_ and *I*_NaP_) working at different timescales and modulated by the presence of noise. In the next section, we show how these observations are connected to oscillations.

### *G*_NaP_, *G*_KDR_ and *G*_h_ determine phase locking of SS and BS to theta and gamma input respectively

It has been demonstrated that this neuron model can reproduce coexisting SS and BS phase-locked to gamma and theta oscillations, respectively, and that these frequency preference (resonance) of SS and BS is interleaved. The model is derived from [1, 15] and has its conductance fitted by obtaining values that generate voltages that match experimental voltage imaging. In this paper, we modified the model for a similar fitting (see parameters in Methods). Based on the observations above for changing ionic conductances and the theta and gamma relationship to BS and SS, we question the ability of the ionic channels to modulate the phase-locking of the neuron model to these frequencies.

Since most of CA1 neuron inputs come with a mixture of theta and gamma frequencies, it is only natural to study the case where both are present. Thus, we injected both theta and gamma simultaneously into the neuron model (Fig 4A). As observed, the ionic currents embedded in the neuron can generate different phase-locking (Fig 4B). Low values of *G*_NaP_ are more prone to BS at theta frequencies (Fig 4C1, Cohen’s d = 1.66; Fig 4C2, Cohen’s d = 2.77). Increasing the values of *G*_NaP_ removes this phase locking with the generation of more SS (Fig 4C3). In contrast, *G*_KDR_ had the opposite behavior. We see only SS for low values of *G*_KDR_ (Fig 4D1), BS at gamma for intermediate values of *G*_KDR_ and finally BS at theta is generated for high *G*_KDR_ (Fig 4D2, Cohen’s d =0.54; Fig 4D3, Cohen’s d = 2.41). Interestingly, only low and high values of *G*_h_ promoted a difference at the gamma frequencies (Fig 4E1, Cohen’s d = 0.68; Fig 4E3, Cohen’s d = -0.26).

**Fig. 4.**
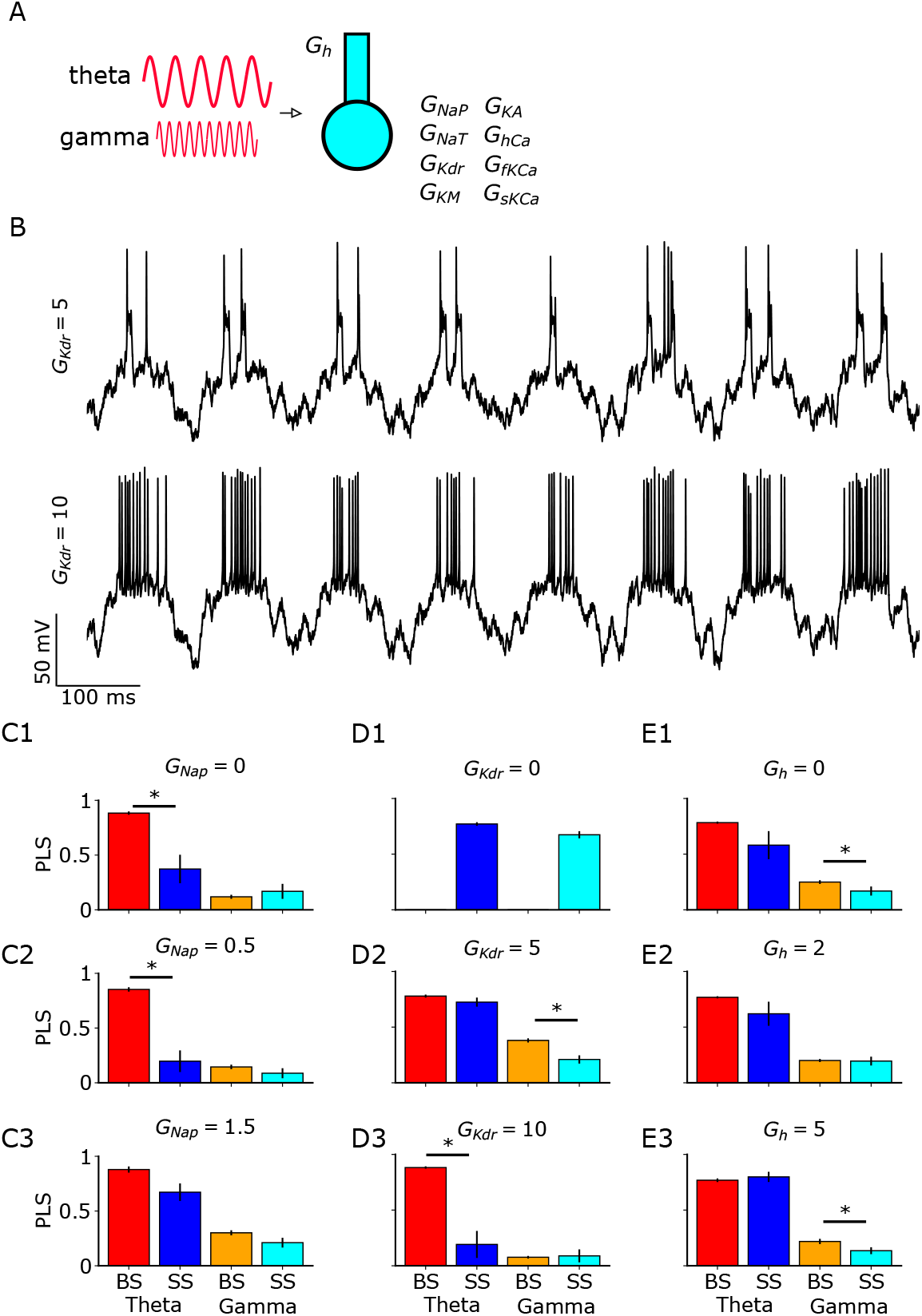
*G*_NaP_, *G*_KDR_ and *G*_h_ determine phase locking statistics (PLS) of SS and BS to theta and gamma input. Default values for fixed conductance while varying the other are *G*_NaP_ = 0.93, *G*_KDR_ = 5.86, and *G*_h_ = 8.46. **A** Sketch of CA1 model with somatic and dendritic compartments receiving theta and gamma oscillatory current stimulation. **B** Examples of voltage traces with SS behavior transitioning to a BS behavior (compare top to bottom). **C1** Phase locking statistic (PLS) to theta and gamma input separated into SS and BS with varying *G*_NaP_ = 0 (asterisk has Cohen’s d 1.66), **C2** when *G*_NaP_ = 0.5 (asterisk has Cohen’s d 2.77) and **C3** when *G*_NaP_ = 1.5. **D1** Phase locking statistic (PLS) with varying *G*_KDR_ = 0, **D2** when *G*_KDR_ = 5 (asterisk has Cohen’s d 0.54) and **D3** when *G*_KDR_ = 10 (asterisk has Cohen’s d 2.41). **E1** Phase locking statistic (PLS) with varying *G*_h_ = 0 (asterisk has Cohen’s d 0.68), **E2** when *G*_h_ = 2 and **E3** *G*_h_ = 5 (asterisk has Cohen’s d -0.26). Each bar is the result of 10 different simulations. t-test with 0.05 level of significance (asterisk).

Taken together, these results show that *G*_NaP_, *G*_KDR_, and *G*_h_ modulate the coexistence of BS and SS phase locked to theta and gamma differently: low values of *G*_NaP_ (Fig 4C1) and high values of *G*_KDR_ (Fig 4D3) lock BS to theta frequencies. *G*_h_ has little effect on theta frequencies and affects gamma frequencies instead. Furthermore, low values of *G*_KDR_ lock SS to gamma frequencies (Fig 4D1).

As previously observed for the remaining ionic currents, we also investigated the separate role of each ionic current in the model and how it would shape the emergence of SS or BS. This would mean either performing a sensitivity analysis or fixing all other parameters and further examining the contribution of all conductances (*G*_KA_, *G*_fKCa_, *G*_KDR_, *G*_KM_, *G*_NaP_, and *G*_sKCa_) to firing frequency and its dependence on input frequency for each pattern (SS and BS). Those results are presented in Fig 5 where heatmaps of firing rates (SS and BS) are displayed as conductance values vary. Our results indicate that calcium-dependent potassium currents (*G*_fKCa_ and *G*_sKCa_) do not influence the firing properties of SS and BS. In contrast, *G*_KA_, *G*_KDR_, *G*_KM_, and *G*_NaP_ exhibit significant effects. Notably, all potassium conductances follow a consistent pattern: increasing their values reduces the input frequency at which SS and BS reach maximum firing. Conversely, *G*_NaP_ shows the opposite effect.

**Fig 5.**
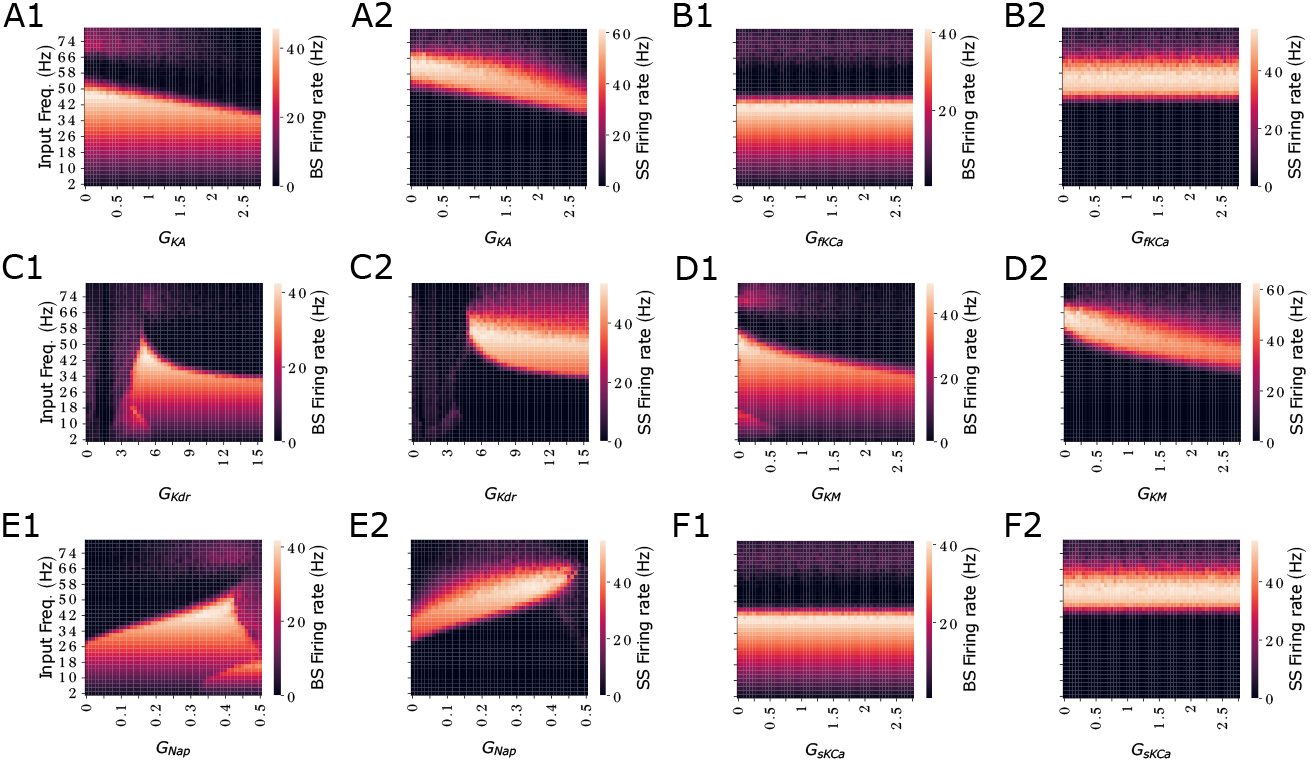
*G*_KA_, *G*_KDR_, *G*_KM_, and *G*_NaP_ determine firing rate of SS and BS. Heatmaps of firing rate for all the conductances for different input frequencies for each pattern (SS and BS). **A1** BS for *G*_KA_ **A2** SS for *G*_KA_. **B1** BS for *G*_fKCa_ **B2** SS for *G*_fKCa_. **C1** BS for *G*_KDR_ **C2** SS for *G*_KDR_. **D1** BS for *G*_KM_ **D2** SS for *G*_KM_. **E1** BS for *G*_NaP_. **E2** SS for *G*_NaP._ **F1** BS for *G*_sKCa_. **F2** SS for *G*_sKCa_.

### Quiet intervals determine the probability of bursting or single spiking

The *in vivo* activity of CA1 neurons is characterized by subthreshold oscillations that follows the theta and gamma synaptic inputs. During these oscillations, spiking activity is observed at the depolarization phase and between them it is possible to observe silence intervals at hyperpolarized phase. During the depolarization, neurons can fire SS or BS. In this section, we investigate whether the silence intervals modulate the probability of having SS or BS.

According to our observations in Fig 6, high probability of BS occurs for long intervals of silence (above 100 ms) in spite of variation in the values of the conductances *G*_NaP_, *G*_KDR_, and *G*_h_ (Fig 6D1, 6E1 and 6F1). Interestingly, we observed that interevent intervals (ISI) are fractured and distributed in more than one position in the diagram, and that different peaks show up mostly for large *G*_NaP_ or low *G*_KDR_ (Fig 6A3 and 6B1). We also observed that high probability of SS occurs for short intervals of silence (below 60 ms) for low *G*_NaP_, high *G*_KDR_ and regardless of the values of *G*_h_ (Fig 6D2, 6E2 and 6F2). Moreover, we observed high probability of SS occurring for long intervals (above 100 ms) only when *G*_NaP_ is high or when *G*_KDR_ is low (Fig 6D2 and 6E2, respectively). In summary, we found that the probability of BS is robust to variation of the ionic conductances and mostly depends on long silent periods above 100 ms (i.e. when theta input is present). In contrast, SS is influenced by ionic conductances and can occur at short or long intervals.

**Figure 6.**
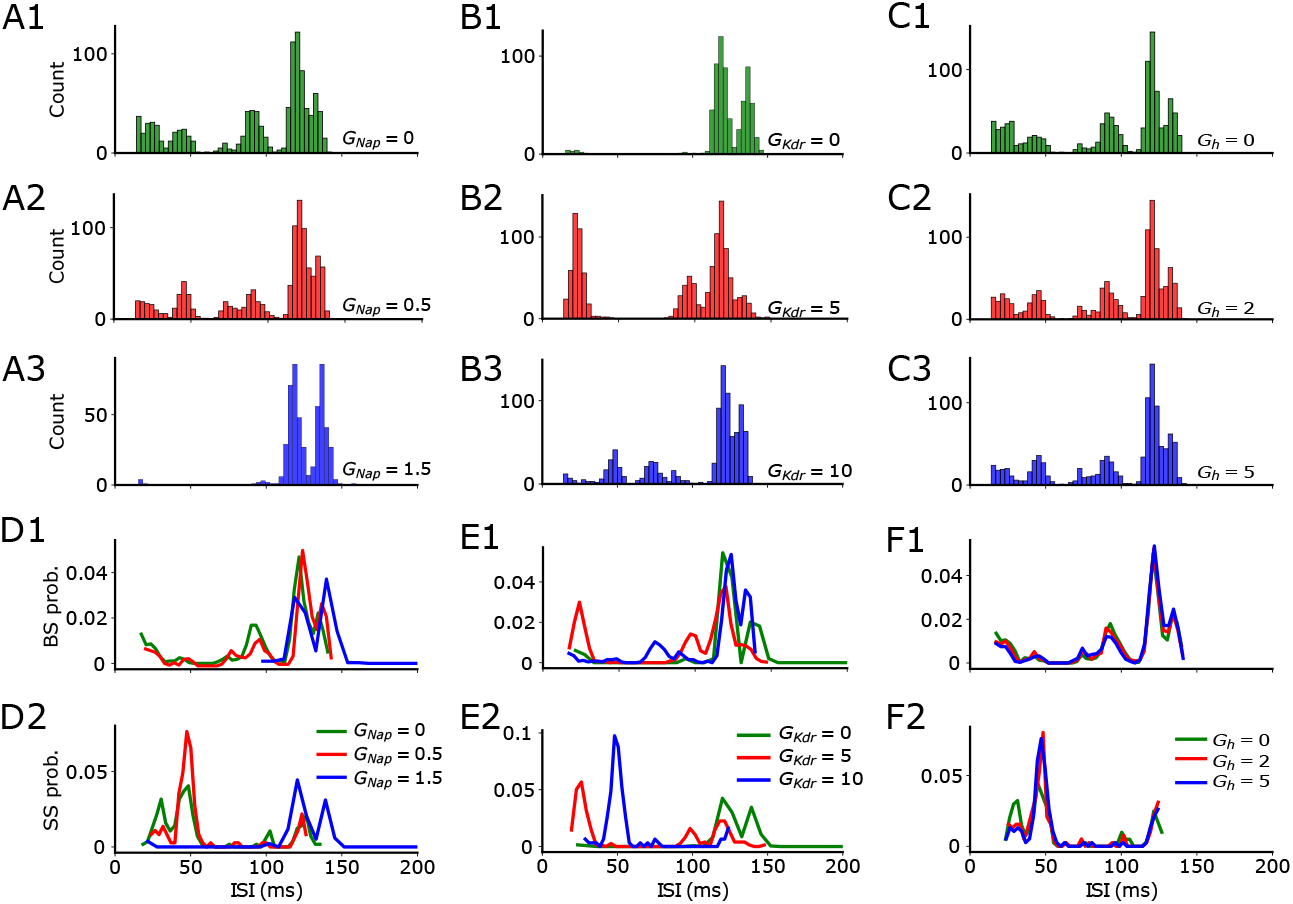
Quiet intervals determine burst and single spikes. Default values for fixed conductance while varying the other are *G*_NaP_ = 0.93, *G*_KDR_ = 5.86, and *G*_h_ = 8.46. **A1-A3** Histogram of interevent intervals (ISI) with varying *G*_NaP_. **B1-B3** Histogram of interevent intervals (ISI) with varying *G*_KDR_. **C1-C3** Histogram of interevent intervals (ISI) with varying *G*_h_. **D1, E1 and F1** BS and **D2, E2 and F2** SS probabilities for varying *G*_NaP_, *G*_KDR_, and *G*_h_, respectively. Colors mark the magnitude of ionic currents (green = low, red = medium, blue = high).

To determine whether the spiking probability is the sum of independent effects of theta and gamma stimulation, or the result of the interaction between both inputs, we calculated the spiking probability when only theta or gamma inputs were used (Fig 7). As expected, when only gamma input is present, we observed that the probability of BS or SS is always larger at short intervals of silence (< 50 ms) (Fig 7A1-7A6). In contrast, when only theta input is present, we observed that the probability of BS is larger for long intervals of silence (> 100 ms) (Fig 7B1-7B3). For SS probability when only theta input is present, we observed the same tendency found above, where high probability of SS occurs for short intervals of silence below 60 ms for low *G*_NaP_, high *G*_KDR_ and regardless the values of *G*_h_, and high probability of SS occurs for long intervals above 100 ms only when *G*_NaP_ is high or when *G*_KDR_ is low (Fig 7B4-7B6).

**Figure 7.**
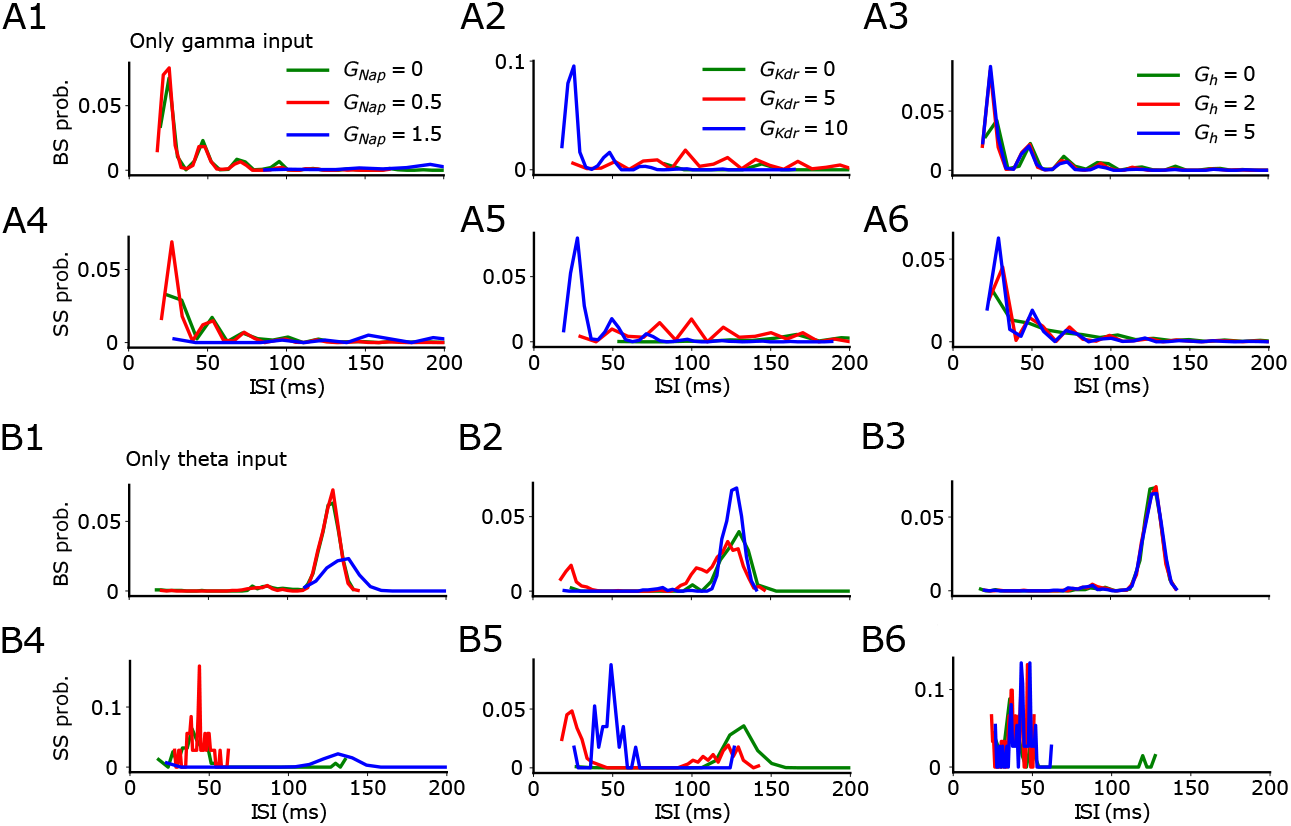
Burst probability follows the longest quiet intervals according to the slowest oscillatory input. Default values for fixed conductance while varying the other are *G*_NaP_ = 0.93, *G*_KDR_ = 5.86, and *G*_h_ = 8.46. **A1-A3** BS and **A4-A6** SS probabilities with only gamma input. **B1-B3** BS and **B4-B6** SS probabilities with only theta input. Colors mark the magnitude of ionic currents (green = low, red = medium, blue = high).

In summary, we observed that when the input is only at gamma frequencies, the spiking probability is largest for short silent intervals for both SS and BS. Opposite to that, when the input is only oscillating at theta, BS is more likely to occur at long intervals and SS at short intervals. Interestingly, when both theta and gamma are present, the probabilities behave as if only theta was present, i.e., the gamma component is mostly disregarded even though the stimulation contains both theta and gamma (Fig 7). These results suggest that BS probability follows the input frequency with the longest intervals of silence, i.e. it follows theta even in the presence of gamma, but it follows gamma in the absence of any other slowest input. Meanwhile, SS would preferentially occur at short intervals.

### Voltage imaging burst probability follows the longest quiet intervals

Despite the difficulties in reconciling our computational results with voltage-imaging recordings, where frequency-dependent stimulation is typically absent, it is possible to compare the bursting probability following the longest quiet intervals with spontaneous recordings. Thus, we analyzed voltage imaging traces of 7 neurons (20 s long) obtained from a public repository [3]. We used a difference source than [1] who were using a general synapsin promoter, hence that was not specific for pyramidal neurons. For our purposes, we chose the dataset available at [3] because the neurons are identified as pyramidal neurons using the CamKII promoter. Fluorescence images were collected from CA1 pyramidal cells and somatostatin expressing interneurons located in the dorsal CA1 area at depths ∼50 to 100 μm (Fig 8A). LED epifluorescence illumination allowed the collection of the voltage indicator fluorescence by a high-speed sCMOS camera at a rate acquisition of 600 Hz. We observed waveform distortions in the voltage imaging traces due to low pass filtering from the slower voltage indicator kinetics relative to the action potential. While this limits the technique, it does not affect our results, as our focus is on peak detection rather than spike kinetics. Photobleaching was subtracted by fitting a single exponential decay. We selected the traces from pyramidal cells based on their high signal-to-noise ratio and high frequency of events suggesting good quality of recordings (Fig 8B). These criteria allowed us to properly detect and classify events as SS and BS as done in the computational simulations (Fig 8C). We calculated the probability of finding SS or BS for a certain ISI (Fig 8D). We performed a BS and SS comparison statistical comparison between the set of ISIs falling in the 3 areas marked for low ISI (100 to 150 ms), intermediate ISI (150 to 250 ms), and long ISI (400 to 500 ms).

**Figure 8.**
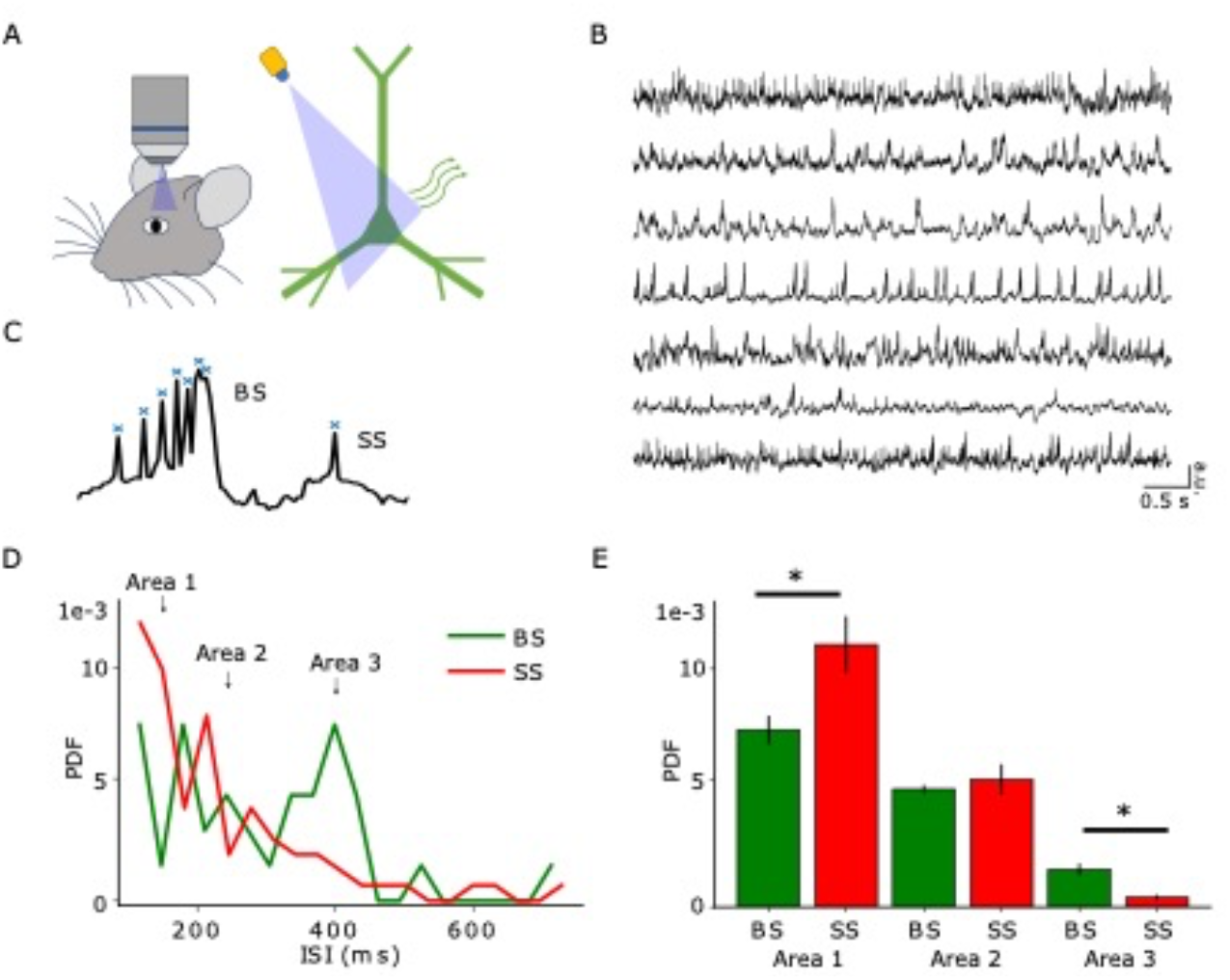
Voltage imaging burst probability follows the longest quiet intervals. Data from [3]. **A** Sketch of the voltage-imaging experiment. **B** Sample traces that were used for this analysis (right, n = 7). **C** Sample trace showing spike detection algorithm prior to division into BS and SS. **D** Interevent interval (ISI) probability density function of BS and SS (n = 7), and 3 areas marked for low ISI (100 to 150 ms), intermediate ISI (150 to 250 ms), and long ISI (400 to 500 ms). **E** From the marked areas, bar plots with averaged values in each ISI trace (n = 7). t-test perfomed with the data in the particular area that delivers 0.05 level of significance (asterisk). Cohen’s d = - 1.02 and p=0.03 for Area 1, Cohen’s d = -0.24 and p=0.68 for Area 2, and Cohen’s d = 1.54 and p=0.02 for Area 3.

The results showed that SS is more likely at low ISIs, while BS is more likely at long ISIs. Observations of these probabilities across all traces (Fig. 8D) and statistical significance from the traces and areas (Fig. 8E) confirm this. The voltage-imaging recordings agree with our simulation observations, where burst probability follows the longest quiet intervals (Figs 6 and 7). An independent study observed this phenomenon in behaving rats, focusing on CA1 pyramidal cell burst activity from electrodes that were surgically implanted within the brain [9].

## Discussion

In this paper, we investigated the role of ionic conductances in causing suprathreshold resonance for SS and BS in a CA1 pyramidal neuron model. When studying neuronal resonance, it is important to assess input-output properties and combinations of intrinsic properties [35, 53, 54, 55, 37, 56]. Our focus lay on specific input and output parameters, with emphasis on oscillatory theta (3-12 Hz) and gamma (30-100 Hz) frequencies for inputs, and SS and BS for outputs. In particular, we systematically assessed these quantitates separately and identified interleaved SS and BS resonance that does not match in frequency nor amplitude, but it is modulated by the ionic currents. To the best of our knowledge, this is the first paper showing the ability of a single neuronal voltage trace to carry a double-coded spiking resonance, where single spikes and burst were separated and shown to have different frequency preferences: interleaved resonance. We have also observed that the burst probability is associated with longer periods of silence and confirmed this observation through the analysis of in vivo voltage-imaging data from CA1 pyramidal neurons.

More specifically, in terms of ionic currents, we found opposite effects of persistent sodium and delayed rectifier potassium currents. Regarding the ionic conductances, we investigate how *G*_NaP_ and *G*_KDR_ ionic currents affect the emergence of SS to BS. Accordingly, whereas BS is observed for large *G*_NaP_ or low *G*_KDR_, SS is observed for low *G*_NaP_ or large *G*_KDR_. We found that under conditions that allow spiking resonance, BS occurs for theta while SS for gamma. Furthermore, at low input frequencies (theta), the firing rate response is higher when *G*_NaP_ is low and *G*_KDR_ is high due to weak low-pass filtering. Conversely, at high input frequencies (gamma), the firing rate response is lower for the same conditions (low *G*_NaP_ and high *G*_KDR_) due to strong filtering. This different strength of filtering is a consequence of the modulation of intrinsic conductances (i.e. *G*_NaP_ and *G*_KDR_). Both conductances cooperate with each other to set the neuronal filtering at different levels for the respective input frequency (i.e. theta and gamma). One consequence of these results is that changes in expression levels in these conductances could allow shifting of the strength of the output for a certain input frequency. As a result, the neuron can facilitate the transfer of low or high frequency inputs according to the expression of its ionic conductances [57, 58, 59].

Future studies should investigate this phenomenon from the point of view of other ionic currents, in particular *G*_KM_ which seems to promote modulation of the bursting behavior (Fig S1) but have not been studied in this work. Other currents such as the calcium-dependent potassium seem to have a minor effect. In addition, we acknowledge that the amplitude current values used in this work are larger than what one would expect experimentally, which is an artifact of the fact that the potassium reversal potential was established at -90 mV. Therefore, values can go as low as -90 mV and around it. However, we have tested smaller amplitude currents (reduced by 2-fold), which cause peak-to-peak membrane potential oscillations of less than 20 mV. This produced similar qualitative results with different quantitative values, such as shifting the resonance frequency (see Fig. S6).

We found that the interplay between ionic currents and oscillatory input frequencies determines the output strength of the firing rate. Specifically, we found that decreasing *G*_NaP_ increases the firing at theta frequencies and decreases the firing at gamma. Again, *G*_KDR_ changes had opposite results. This suggests that CA1 neurons might be able to tune their output preferences by downregulating or upregulating the expression of their intrinsic conductances [60]. It also suggests that pharmacological interactions targeting these ionic channels have the potential to affect oscillations and BS behavior [61, 62].

We previously found that BS and theta frequencies are preferentially linked to the animal entering a place field, while SS and gamma frequencies are linked to the animal leaving a place field [1]. Thus, BS and SS can be considered coding mechanisms for this type of behavior. Our simulations demonstrate that different oscillatory frequencies can enhance (resonate with) BS and SS separately in a complex frequency-dependent manner within this double-coded signal. This indicates a higher-order interaction where the neuron does not need to code either BS or SS exclusively but can encode both jointly. It would be interesting to find ways in which one resonance is kept fixed while the other is varied. This would prove whether they are dependent or independent.

An interesting finding of our work was that ionic currents embedded in the neuron can generate different phase-locking. These results showed that *G*_NaP_, *G*_KDR_, and *G*_h_ modulate the coexistence of BS and SS phase locked to theta and gamma differently: while low values of *G*_NaP_ generate BS at theta, it requires high values of *G*_KDR_ to generate BS at theta. *G*_h_ has little effect on theta frequencies and affects gamma frequencies instead (Fig 4 E1-E3). *I*_h_ current modulation of up/down states is still under investigation [63], but clearly a target of a diverse range of neuromodulators [64, 65].

Unexpectedly, we found that the likelihood of BS remains consistent despite changes in ionic conductances and primarily relies on extended silent periods [9], meanwhile SS is influenced by ionic conductances changes and can occur at any intervals. This BS behavior can be explained by the degeneracy of ionic conductances. In these circumstances, multiple combinations of different conductance values, even with significant heterogeneity, can produce the same BS behavior in the neuron model [60, 66]. Another possible interpretation is that the various conductances within the neuron model (such as transient sodium, muscarinic-sensitive potassium, A-type potassium, high-threshold calcium, fast calcium-activated potassium, and slow calcium-activated potassium currents) might respond differently to prolonged hyperpolarization, primarily inducing a state conducive to bursting rather than single spiking. Conversely, shorter hyperpolarizations could result in bistability, where both bursting and single spiking scenarios are plausible [67, 68].

The probability of spiking is influenced by the states of the conductances (i.e., the levels of activation/inactivation) before the onset of depolarization [69]. For the conductances examined in this paper, it is well known that during extended hyperpolarization (such as under the stimulation of theta frequency oscillations), *G*_KDR_ and G_NaP_ remain inactive. However, during the initial phase of depolarization, both *G*_KDR_ and *G*_NaP_ are rapidly activated [41], leading to the onset of bursting. Similarly, it has been observed that the interplay between potassium and sodium conductances triggers bursting after extended periods of silence [9, 67]. These findings align with previous research demonstrating that these currents (e.g., *G*_KDR_ and *G*_NaP_) are crucial for generating bursting [15, 70]. Finally, it has been observed in cortical neurons that bursts can trigger switches from intermittent bursting spiking to persistent single spiking activity and vice-versa [71]. These findings bring new insights into the neural coding mechanisms in the CA1 hippocampus.

## Methods

### Computational model

We use a Hodgkin-Huxley model of a CA1 pyramidal neuron described by conductance-based ionic currents [15, 1, 72, 73, 74, 75, 76]. The following differential equation describes how the voltage evolves in time:

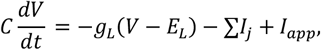

with the currents *I*_j_

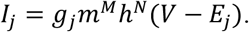

The currents included are *I*NaT the transient sodium (M = 3, N = 1), *I*NaP the persistent sodium (M = 1, N = 0), *I*Kdr the delayed rectifier K^+^ (M = 0, N = 4), *I*KM the muscarinic-sensitive potassium (M = 1, N = 0), *I*KA the A-type K^+^ (M = 3, N = 0), *I*HCa the high-threshold Ca^2+^ (M = 2, N = 0), *I*fKCa the fast Ca^2+^ -activated potassium current (M = 1, N = 0), and *I*sKCa the slow Ca^2+^-activated potassium current (M = 1, N = 0). In addition, when the dendrite is present it contains an *I*h current (M=1, N=0). The reversal potentials are *E*_L_ = −70 mV, *E*_K_ = −90 mV, *E*_Na_ = 55 mV, and *E*_Ca_ = 120 mV. The remaining parameters are *C* = 1 μF/cm^2^, *g*_L_ = 0.04 mS/cm^2^, *g*_NaT_ = 35 mS/cm^2^, *g*_NaP_ =0.9 mS/cm^2^, *g*_Kdr_ = 5.8 mS/cm^2^, *g*_KA_ = 2.8 mS/cm^2^, *g*_h_ =8.46 mS/cm^2^, *g*_KM_ = 2.4 mS/cm^2^, *g*_HCa_ = 0.17 mS/cm^2^, *g*_sKCa_ = 2.8 mS/cm^2,^ *g*_*fKCa*_= 9.05 mS/cm^2^. Throughout the paper, all conductance values will be displayed in mS/cm^2^, but we will omit the units unless necessary for clarity. All currents in the model have units μA/cm^2^, but we will also omit units. Equations for *m* and *n* follow

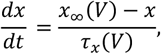

where 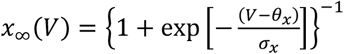. This is the same for all ionic currents except to *I* which has *d* variable governed by calcium following *d*_*∞*_ ([*Ca*^2+^]_*i*_) = {1 + a_*c*_/[*Ca*^2+^]_*i*_}^−1^ and *I*sKCa which has a *q* variable governed by calcium following 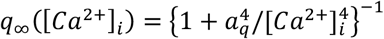.

For the dynamics of the calcium concentration inside the cell [*Ca*^2+^]_*i*_ we assume

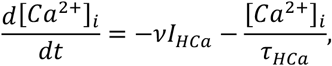

where ν = 0.13 cm^2^/(ms× μA), and τ_*HCa*_ = 13 ms.

The remaining parameters for these equations are *θ*_*m*_= −30 mV, σ_*m*_ = 9.5 mV, *θ*_*h*_ = −45 mV, σ_*h*_ = −7 mV, *θ*_*p*_ = −35 mV, σ_*p*_ = 3 mV, *θ*_*n*_ = −35 mV, σ_*n*_ = 10 mV, *θ*_*nt*_ = −27 mV, σ_*nt*_ = −15 mV, *θ*_*a*_ = −50 mV, σ_*a*_ = 20 mV, *θ*_*b*_ = −80 mV, σ_*b*_ = −6 mV, *θ*_z_ = −39 mV, σ_z_ = 5 mV, *θ*_*r*_ = −20 mV, σ_*r*_ = 19 mV, *θ*_*c*_ = 30 mV, σ_*c*_ = −7 mV as in [15].

For the compartmental modeling, we add axial conductance between *i* and *i+1* following the ohmic current *I*_*ac*_ = *g*_*ac*_ (*V*_*i*_ − *V*_*i*+1_), with *g*_*ac*_ = 0.2 mS/cm^2^ For external currents, we use sinusoidal currents with specific frequencies or combinations thereof, as illustrated in the respective figures. Gamma frequencies are defined as ranging from 30 to 100 Hz, and theta frequencies range from 3 to 12 Hz. In our simulations, we use either 40 Hz for gamma or 10 Hz for theta. We chose sinusoidal current injections over conductance-based synapses because our previous work [38] showed that cells exhibit impedance peaks (Z-resonance) and voltage envelope resonance in response to sinusoidal inputs but not to square-wave or synaptic-like inputs. This applies to both conductance- and current-based inputs. Thus, we used sinusoidal stimulation, consistent with prior experimental approaches [1].

A complete description of the model can be found in the table below.

**Table 1.** Summary of parameters and variables used to build the computational model.

| Ionic currents | Time constant, ms | Parameters |
| --- | --- | --- |
| $I_{\text{NaT}}(V, h) = g_{\text{NaT}} m_\infty^3(V) h(V - V_{\text{Na}})$ | $\tau_h = 0.1 + 0.75 \times \left\{1 + \exp\left[-\frac{V - \theta_{ht}}{\sigma_{ht}}\right]\right\}^{-1}$<br>$m = m_\infty$ | $\theta_m = -30 \text{ mV}$,<br>$\sigma_m = 9.5 \text{ mV}$,<br>$\theta_h = -45 \text{ mV}$,<br>$\sigma_h = -7 \text{ mV}$ |
| $I_{\text{NaP}}(V) = g_{\text{NaP}} p_\infty(V)(V - V_{\text{Na}})$ | $p = p_\infty$ | $\theta_p = -35 \text{ mV}$,<br>$\sigma_p = 3 \text{ mV}$ |
| $I_{\text{Kdr}}(V, n) = g_{\text{Kdr}} n^4(V - V_K)$ | $\tau_n = 0.1 + 0.5 \times \left\{1 + \exp\left[-\frac{V - \theta_{nt}}{\sigma_{nt}}\right]\right\}^{-1}$ | $\theta_n = -35 \text{ mV}$,<br>$\sigma_n = 10 \text{ mV}$ |
| $I_{\text{KM}}(V, z) = g_{\text{KM}} z(V - V_K)$ | $\tau_z = 75$ | $\theta_z = -39 \text{ mV}$,<br>$\sigma_z = 5 \text{ mV}$ |
| $I_{\text{KA}}(V, b) = g_{\text{KA}} a^3(V) b(V - V_K)$ | $a = a_\infty$<br>$\tau_b = 15$ | $\theta_a = -50 \text{ mV}$,<br>$\sigma_a = 20 \text{ mV}$,<br>$\theta_b = -80 \text{ mV}$,<br>$\sigma_b = 15 \text{ mV}$ |
| $I_h(V, ) = g_h t(V)(V - V_K)$ | $\tau_t = \{\exp[-17.9 + 0.116V] + \exp[-1.84 + 0.09V]\}^{-1} + 0.1$ | $\theta_a = -84 \text{ mV}$,<br>$\sigma_a = 10.2 \text{ mV}$ |
| $I_{\text{HCa}}(V, r) = g_{\text{HCa}} r^2(V - V_{\text{Ca}})$ | $\tau_r = 1$ | $\theta_r = -20 \text{ mV}$,<br>$\sigma_r = 19 \text{ mV}$ |
| $I_{\text{fKCa}}(V, c) = g_{\text{fKCa}} d_\infty([Ca^{2+}]_i) c(V - V_K)$ | $\tau_c = 2$ | $\theta_c = 30 \text{ mV}$, $\sigma_c = -7 \text{ mV}$, $a_c = 6 \text{ mV}$ |
| $I_{sKCa}(V, q) = g_{sKCa}q(V - V_K)$ | $\tau_q = 450$ | $a_q = 2 \text{ mV}$ |
| **Current between compartments** |  |  |
| $I_{ac} = g_{ac}(V_i - V_{i+1})$ | | $g_{ac} = 0.2$ |
| **External current** |  |  |
| $I_{sin} = I_\theta + I_\gamma$ | $I_\theta$ is a sinusoidal 10Hz current and $I_\gamma$ is 40Hz, with amplitudes 1 $\mu\text{A}/\text{cm}^2$ and 2 $\mu\text{A}/\text{cm}^2$, respectively. | |
| $I_{app}$ | Amplitude 1 $\mu\text{A}/\text{cm}^2$ | |
| **External Noise** |  |  |
| $I_\eta = \sqrt{2D}\xi(t)$ | The function $\xi(t)$ is a zero-mean white Gaussian noise with $\langle \xi(t)\xi(t') \rangle = \delta(t - t')$. | $D = 9.6 \times 10^{-6}$ |

The code for this model is available at https://github.com/rodrigo-pena-lab/interleaved_resonance.

### Experimental recordings

To compare the model results with experimentally recorded in vivo voltage-imaging using genetically encoded voltage indicator (acemNeon2) in awake moving mice, we used freely available data from [3] in https://tinyurl.com/kannanEtAlScience2022 (last accessed in June 2024). The recordings are 20 seconds long and the data for this paper contain 7 CA1 pyramidal cells from depths ranging from approximately 50 to 100 μm. Image segmentation was performed using custom-written Matlab scripts as used in previous publications [45, 77, 78, 79]. CA1 pyramidal cells were identified by their expression of the negative polarity relationship of fluorescence emission to the membrane voltage as expected from the voltage indicator (acemNeon2), whereas somatostatin interneurons were identified by the positive polarity that expressed a different voltage indicator (pACe). The voltage indicator was injected in the entorhinal cortex (EC) and expressed retrogradely in CA1, thus, all CA1 pyramidal neurons reported here are projecting to EC. Data acquisition was performed using LED epifluorescence illumination, and the voltage indicator fluorescence was recorded at a rate of 600 Hz with a high-speed sCMOS camera. To correct for photobleaching, we applied a single exponential decay fit and subtracted it from the data. We specifically selected traces from pyramidal cells that exhibited a high signal-to-noise ratio and a high frequency of events, which indicated good recording quality.

### Firing rate and coefficient of variation

We compute the firing rate as the sum of spikes divided by the time window considered. Unless stated otherwise, we adopted 4 s in each simulation. The inter-spike intervals (ISI) are taken as the intervals between spikes in the trace. From the ISI we define the coefficient of variation (CV) which points to burstiness in the traces for CV>2, and is defined as the standard deviation over the mean of the ISI:

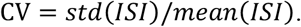

We also divide intervals between single-spikes and bursts accordingly. ISI probabilities are calculated using the same criteria for separating ISIs from single spikes and bursts. The intervals are measured from the last spike to the next single spike or burst spike, regardless of whether the last spike was part of a single spike or a burst.

### Burst detection

Following [4] that showed through intracellular studies that individual spikes with complex spikes can have ISIs of more than 10 ms, we adopted a 14ms ISI time window to separate single-spikes from bursts. At least two spikes are necessary for this analysis, otherwise the spike-train is disregarded. As for the detected burst, we separate data from the first single spike only to account for the number of bursts and not the intraburst frequency.

### Spike phase-locking computation

We quantify the relative phase of a spike to an oscillation using the phase-locking value (PLV):

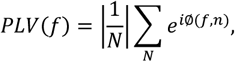

with *f* the frequency and *N* the number of spikes [80]. For theta oscillations we assume 3-12 Hz and for gamma oscillations we assume 30-100Hz. The phase *ϕ* was obtained with the help of the Hilbert transform. The phase-locking statistics (PLS) accounts for uneven size in the data between bursting and single-spikes [81, 82]:

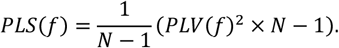

### Resonance frequencies and amplitudes

We compare resonance frequencies and their amplitudes using the following equations: Δ*f* = *f*_*SS*_ − *f*_*BS*_ and Δ*A* = *A*_*SS*_ − *A*_*BS*_, where f_SS_ is the resonance frequency of single spikes and f_BS_ is the resonance frequency of bursts (in Hz). Also, A_SS_ is the firing frequency at the resonance peak (also known as firing rate amplitude) of single spikes and A_BS_ is the firing frequency at the resonance peak of bursts (in Hz).

## Data availability

Data will be made available on request.

## Supporting information

**Fig S1.**
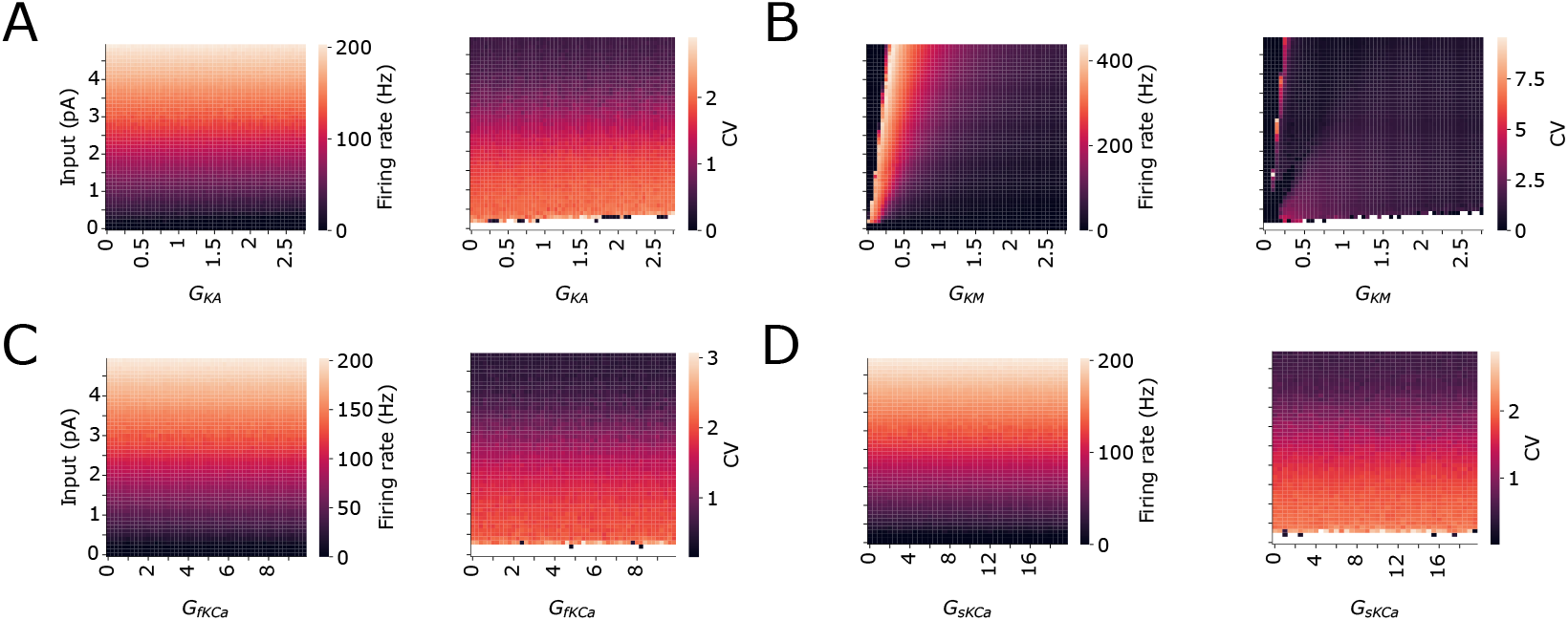
Ionic currents effect on neuronal response during sustained depolarization. Firing rate and coefficient of variation (CV), a measure the activity of the neuron and the presence of bursts, displayed as heatmaps spanned over different ionic current variation. **A** *G*_KA_ vs. Input current. **B** *G*_KM_ vs. Input current. **C** *G*_*f*KCa_ vs. Input current. **D** *G*_*s*KCa_ vs. Input current.

**Fig S2.**
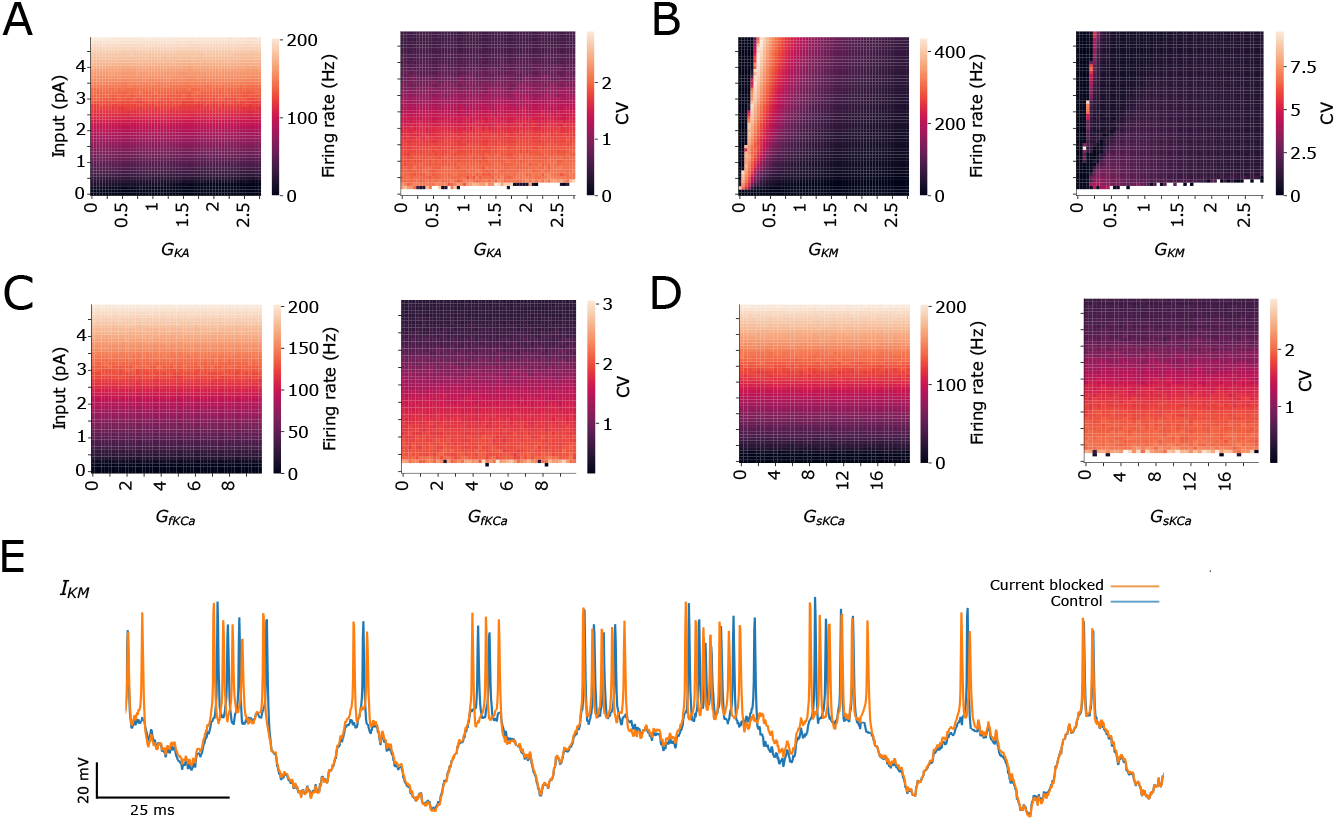
Ionic currents effect on neuronal response during oscillatory input. Firing rate and coefficient of variation (CV), a measure the activity of the neuron and the presence of bursts, displayed as heatmaps spanned over different frequency input variation. **A** *G*_KA_ vs. Input current. **B** *G*_KM_ vs. Input current. **C** *G*_*f*KCa_ vs. Input current. **D** *G*_*s*KCa_ vs. Input current. **E** Voltage traces of control vs after blocking M-type potassium current.

**Fig S3.**
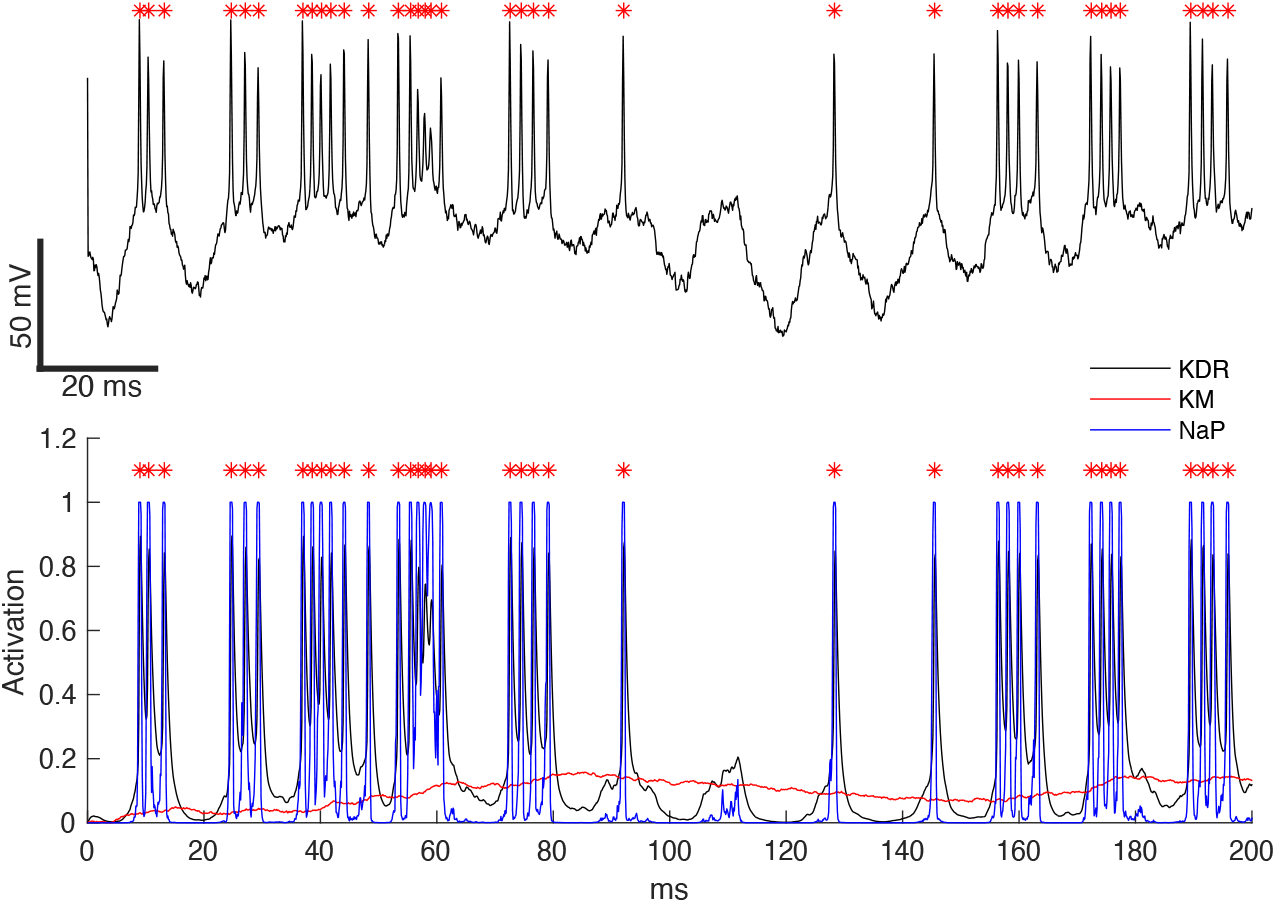
Time evolution of activation variables. **A** Voltage trace showing interleaved single spiking and bursting during oscillatory stimulation. Asterisks represent spike peaks. **B.** Time evolution of activation variables of *I*_KDR_, *I*_KM_ and *I*_NaP_. Note the different temporal scales.

**Fig S4.**
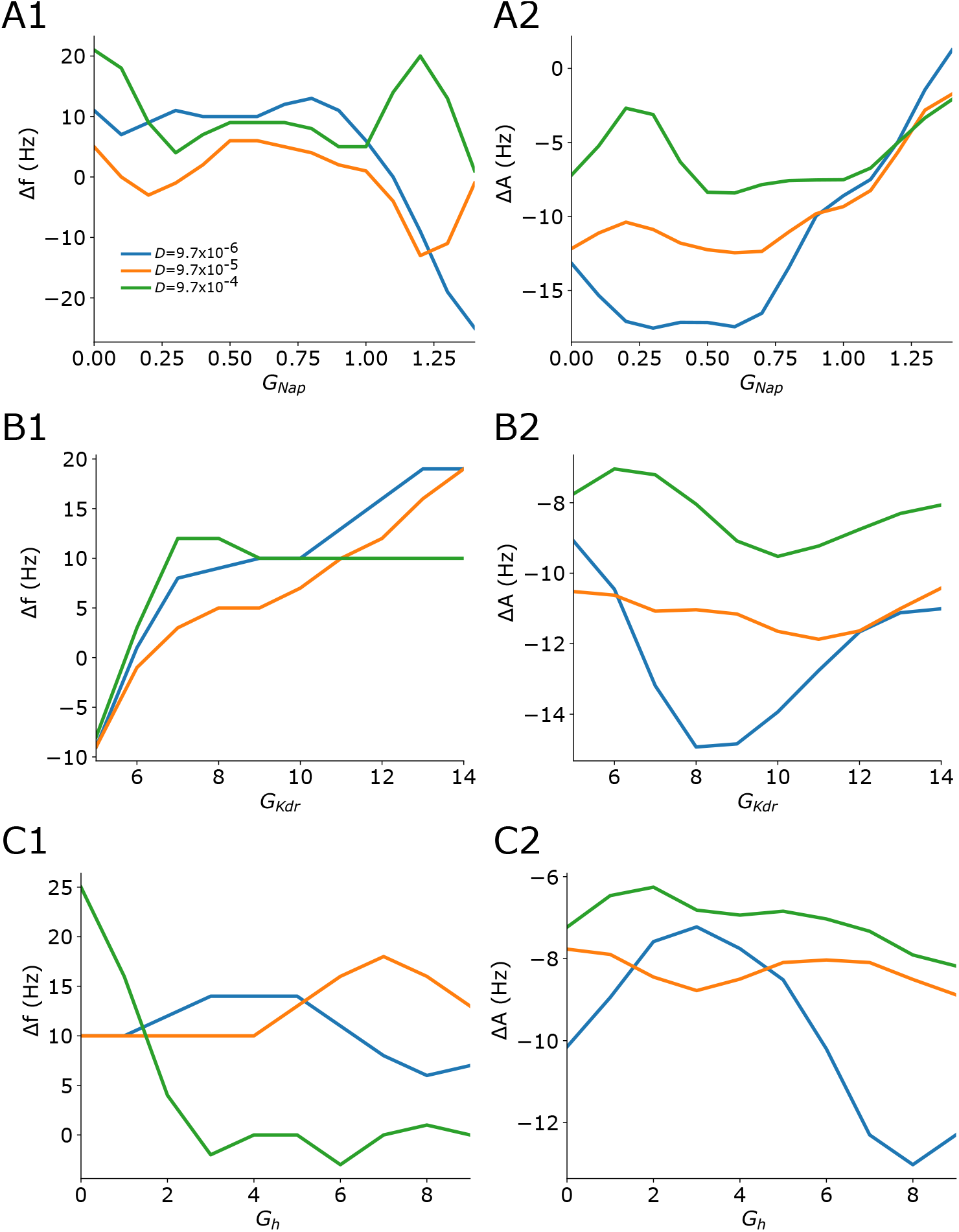
Effect of noise on interleaved resonance. Default values for fixed conductance while varying the other are *G*_NaP_ = 0.93, *G*_KDR_ = 5.86, and *G*_h_ = 8.46. **A1-2** Δ*f* and Δ*A* for varying values of *G*_NaP_. **B1-2** Δ*f* and Δ*A* for varying values of *G*_KDR._ **C1-2** Δ*f* and Δ*A* for varying values of *G*_h_. Colors mark the magnitude of noise (blue = low, orange = medium, green = high).

**Fig S5.**
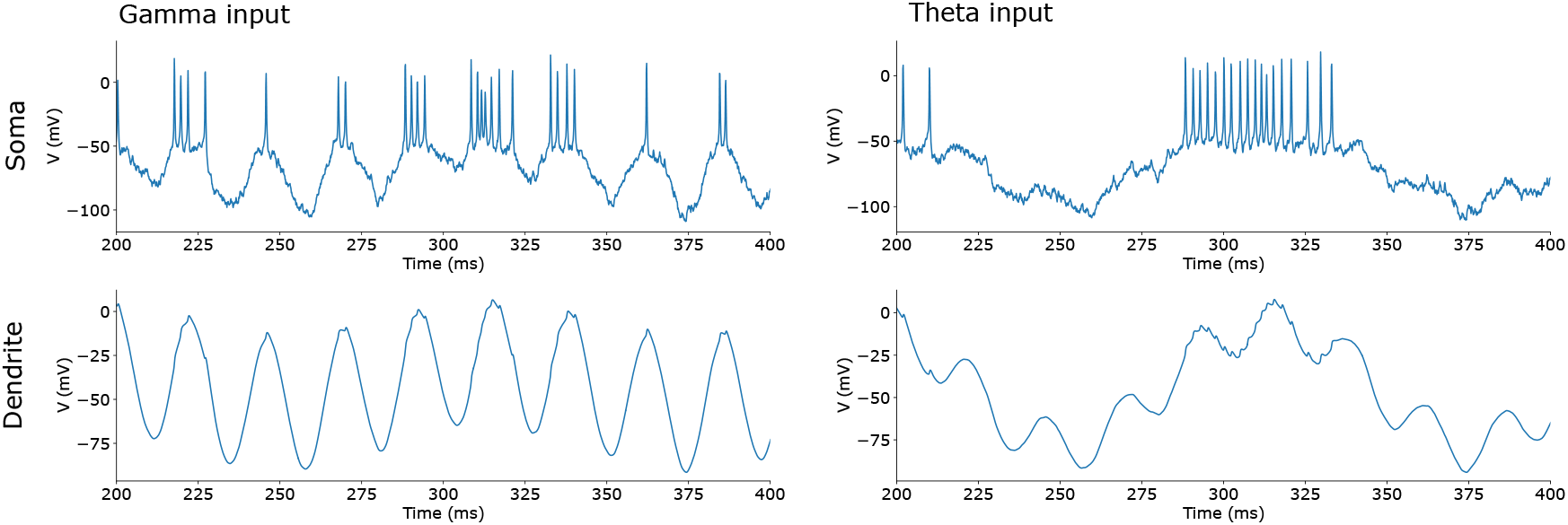
Somatic and dendritic traces under gamma and theta oscillations input. Somatic traces (top) and dendritic traces (bottom).

**Fig S6.**
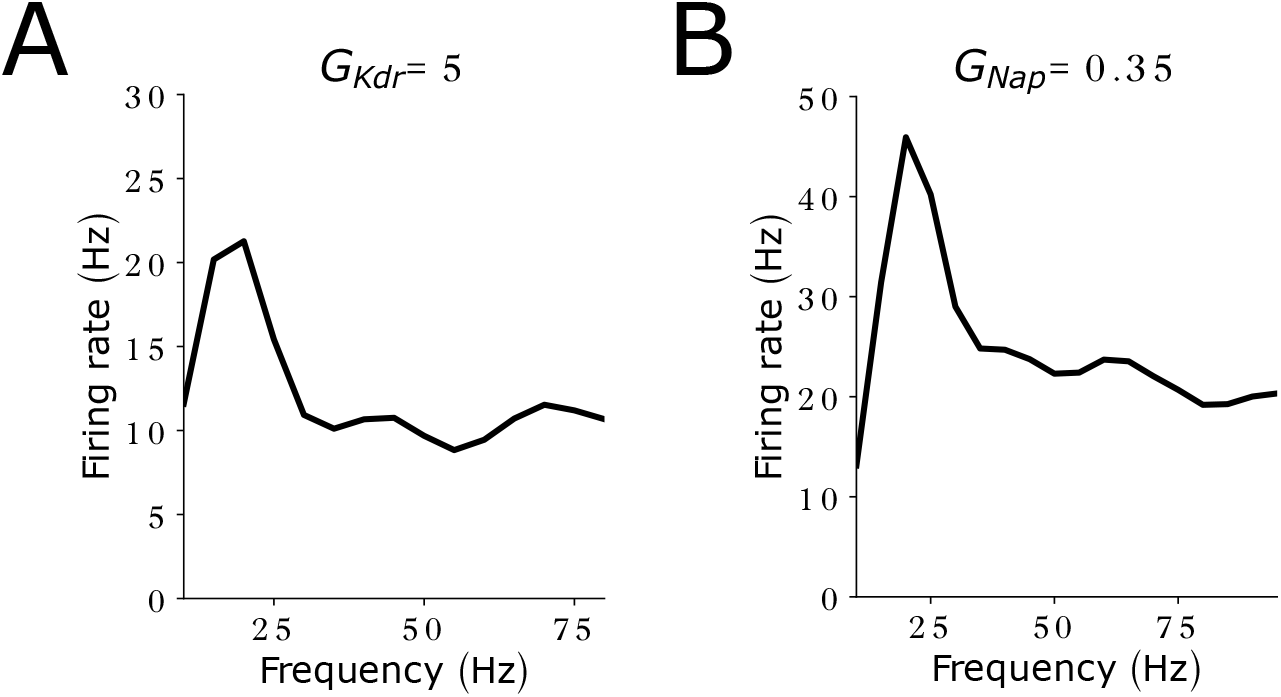
Intermediate *G*_NaP_ and *G*_KDR_ generates firing rate resonance even at low input amplitudes. Same simulation as in Fig 2, but for lower input amplitudes (2x). We have also raised the holding potential by to be closer to threshold for this experiment (offset amplitude: 2 μA/cm^2^**) A** Firing rate while varying stimulation frequency of an oscillatory input for *G*_KDR_ = 5. **B** Firing rate while varying stimulation frequency of an oscillatory input for *G*_NaP_ = 0.35.

